# A nuclear role for the contractile protein troponin I/UNC-27 in regulating muscle aging in *C. elegans*

**DOI:** 10.64898/2026.08.08.743652

**Authors:** Allan Alcolei, Manon Froment, Laurent Molin, Charline Roy, Romain Bulteau, Jean-Louis Bessereau, Florence Solari

## Abstract

Muscle ageing is characterized by evolutionarily conserved subcellular alterations across diverse organisms. In *Caenorhabditis elegans*, the decline in sarcomeric gene expression is among the earliest detectable ageing-associated changes, emerging at the onset of adulthood. To identify causal regulators of muscle ageing in an unbiased manner, we developed a genetic screening strategy that enables visual monitoring of muscle ageing at both cellular and organismal scales. Using this approach, we identified a mutation that delays the age-associated loss of sarcomeric transcripts. Unexpectedly, the mutation maps to the troponin I gene *unc-27*, which encodes a conserved regulator of muscle contraction not previously implicated in gene regulation. The mutation alters a single amino acid within a predicted nuclear localization signal (NLS). We found that multiple NLS motifs mediate the active transport of UNC-27 into muscle nuclei from early adulthood onward. Disruption of UNC-27 nuclear localization preserves sarcomeric gene expression during ageing and delays early hallmarks of muscle decline, including proteostatic imbalance and mitochondrial fragmentation. Transcriptomic analyses further revealed that nuclear UNC-27 selectively regulates the expression of genes encoding structural components of the muscle apparatus in adult animals.

These results support the existence of a homeostatic sarcomere surveillance pathway, in which a structural protein unexpectedly acquires a transcriptional regulatory role in response to age-associated physiological state. The conservation of NLS motifs in mammalian UNC-27 orthologues suggests that this mechanism may be evolutionarily conserved, with potential relevance to human muscle physiology and disease.

## Introduction

Muscle ageing is characterized by progressive alterations in muscle structure, mass, and function, the interrelationship of which remains incompletely understood. Disentangling causal relationships among these parameters is particularly challenging in complex organisms, where longitudinal analyses at subcellular resolution are limited. To overcome these constraints, we used the transparent, short-lived nematode *Caenorhabditis elegans*, a powerful model system that enables longitudinal studies at individual tissue scale in living animals, facilitating the identification of causal links between early molecular events and subsequent functional decline.

Striated body-wall muscles (BWM) in *C. elegans* share several molecular and subcellular features of aging with mammalian cardiomyocytes and skeletal myofibers, including downregulation of sarcomeric genes, impaired proteostasis, mitochondrial dysfunction, and altered calcium handling (Dridi et al., 2022; Gaffney et al., 2018; Mergoud Dit Lamarche et al., 2018). We previously demonstrated that the decline in sarcomeric gene expression is one of the earliest detectable changes in BWM during adulthood (Mergoud Dit Lamarche et al., 2018). More recent studies have extended this observation across species, showing that sarcomeric gene expression decreases with age in *Drosophila*, rodents, and human cardiac and skeletal muscles (Kedlian et al., 2024; Lai et al., 2024; Zhang et al., 2023). While the regulation of sarcomeric gene expression during development has been extensively characterized (for a review, see Bentzinger et al., 2012), the mechanisms governing its regulation during adulthood remain largely unexplored.

In earlier work using a candidate-gene approach, we identified the MADS-box transcription factor UNC-120, the orthologue of serum response factor (SRF), as a key regulator required for muscle maintenance from middle adulthood onward (Mergoud Dit Lamarche et al., 2018). Notably, its role appears evolutionarily conserved, as myofiber-specific inactivation of SRF in mice accelerates muscle ageing (Lahoute et al., 2008; Sakuma et al., 2008). Furthermore, we demonstrated that lifespan regulation and muscle aging can be genetically dissociated (Mergoud Dit Lamarche et al., 2018), strongly suggesting that conventional lifespan-based screens for aging regulators may have overlooked key modulators of muscle aging.

To identify regulators of age-dependent changes in sarcomeric gene expression in an unbiased manner, we designed an innovative genetic strategy based on visual monitoring of muscle aging at individual and cell scales. Following random mutagenesis of the reporter strain, this approach provides a means to identify mutants that maintain sarcomeric gene expression with aging. To our surprise, we identified here the sarcomeric protein troponin I (UNC-27) as a novel regulator of muscle aging. Troponin I is a core component of the troponin complex, which, together with troponin T and troponin C, regulates calcium-dependent muscle contraction. The complex is associated with actin thin filaments *via* tropomyosin: troponin T anchors the complex, troponin C binds calcium, and troponin I inhibits actin– myosin interaction. Upon calcium binding, troponin C undergoes a conformational change that relieves troponin I–mediated inhibition, thereby enabling actin–myosin interaction and muscle contraction. To date, as in mammals, *unc-27* mutations have primarily been reported to affect muscle contraction and the stability of sarcomeric structure (Barnes et al., 2016).

Here, we show that UNC-27 has also a nuclear role in repressing sarcomeric gene expression that is distinct from its canonical role at the sarcomere. UNC-27 nuclear role emerges within a couple of days after the young adult stage and shows that the decrease of muscle-specific transcripts with age is not incidental but under the control of a dynamic process. Collectively, these findings identify a previously unrecognized moonlighting function of UNC-27, which emerges specifically in aging muscle and may be part of an inhibitory feedback regulation between structural proteins and gene expression, offering novel insights into the molecular drivers of muscle aging and their regulation.

## Results

### Identification of a novel *unc-27* mutant that impacts sarcomeric transcripts with aging

To uncover novel regulators of muscle aging, we developed an innovative genetic strategy that enables visual monitoring of age-dependent changes in sarcomeric transcript levels. The *tnt-2* gene was chosen as its transcripts level is one of the most strongly downregulated during early adulthood in *C. elegans* (Mergoud Dit Lamarche et al., 2018). We generated transgenic worms expressing a fluorescent *tnt-2* reporter by inserting a F2A::TagRFP-T::PEST reporter cassette downstream of the endogenous *tnt-2* locus *via* CRISPR/Cas9-mediated genome editing. The F2A sequence mediates ribosomal skipping, while the PEST motif encodes a protein destabilization domain (**Fig. 1A**). The reporter strain shows a pronounced age-dependent reduction in *tnt-2* expression, denoted by the marked decrease in fluorescence observed in 6-day-old worms (**Fig. 1B**). Following random mutagenesis with ethyl methanesulfonate, we conducted a small-scale pilot screen of 1,000 genomes to identify mutants exhibiting delayed downregulation of *tnt-2* expression. Chemically-induced nucleotide changes can lead to various types of mutations providing valuable insights into structure-function relationship as compared to an RNAi screening strategy. Using whole-genome sequencing, genetic mapping, and CRISPR/Cas9-mediated correction of genetic variants (see Methods), we identified a point mutation in *unc-27* resulting in a cysteine-to-arginine substitution at position 24 of the protein (hereafter *unc-27(R24C)*). Introduction of this same mutation by CRISPR/Cas9 in the original reporter strain recapitulated the observed phenotype, thus demonstrating the causal role of this mutation.

**Fig. 1.**
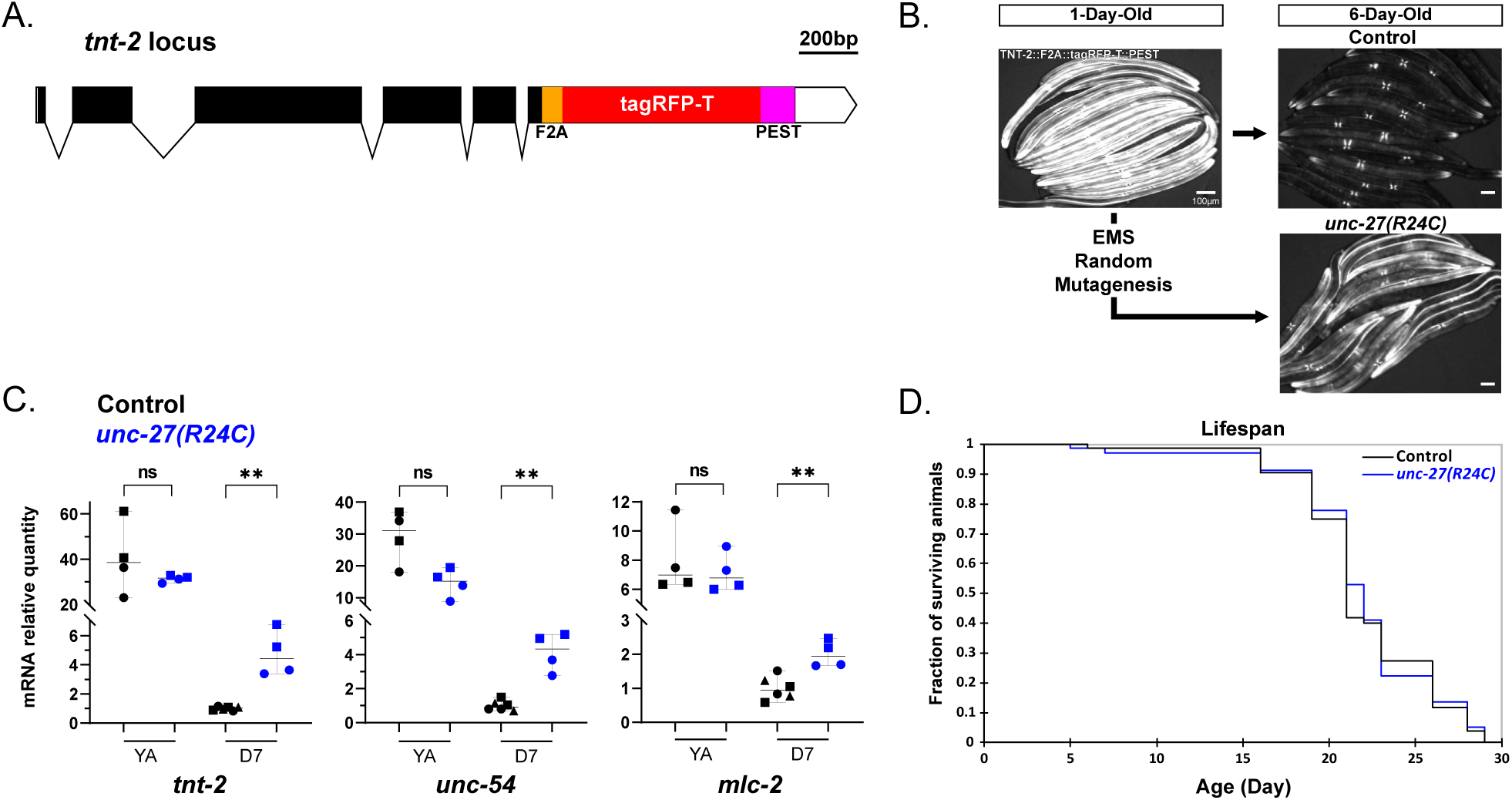
Identification of UNC-27 as a regulator of sarcomeric gene expression during adulthood. (A) Schematic representation of the reporter cassette (F2A::TagRFP-T::PEST) inserted at the endogenous *tnt-2* locus. (B) Representative fluorescence images of *tnt-2* expression reporter in the body-wall muscle (BWM) of wild-type (WT) and *unc-27(R24C)* mutant isolated from the screen at day 1 and day 6 of adulthood. Scale bars, 100 μm. (C) Relative transcript levels of the muscle genes *tnt-2*, *unc-54*, and *mlc-2* in young adults and day-7 adults in control and *unc-27(R24C)* backgrounds. Each dot represents an individual replicate; biological and technical replicates are distinguished by different and identical shapes, respectively. Bars show median ± 95 % confidence interval. Statistical significance was determined by two-sided Mann-Whitney tests; **, *p*<0.01; ns, not significant. (D) Survival curve analysis of control and *unc-27(R24C)* worms with an average lifespan of 22 days (+/-0.5; n=77) and 21,9 days (+/-0.5; n=72) respectively for WT and mutants, Log-rank test *p*-value=0.841.

Quantitative PCR confirmed that the maintenance of fluorescence in aging *unc-27(R24C)* mutants results from increased *tnt-2* transcript levels in 7-day-old adults compared to wild-type animals (**Fig. 1C**). This effect is not restricted to *tnt-2*: *unc-27(R24C)* mutants also maintain elevated transcript levels of other sarcomeric genes that normally decline with age (Mergoud Dit Lamarche et al., 2018), including the myosin heavy and light chains encoded by *unc-54* and *mlc-2*, respectively (**Fig. 1C**). While transcript levels were comparable between wild-type and *unc-27(R24C)* animals at the young adult stage, they were all significantly increased in 7-day-old *unc-27(R24C)* mutants relative to age-matched wild-type animals (**Fig. 1C**). Notably, *unc-27*(*R24C)* mutants exhibit a wild-type lifespan (**Fig. 1D**), suggesting that UNC-27 does not impact the general organismal homeostasis but instead participates in a regulatory mechanism controlling muscle gene expression during aging.

### *unc-27(R24C)* mutants show preserved myofilament organization

Previous studies of *unc-27* mutants demonstrated that UNC-27 is required for both normal locomotion and maintenance of sarcomeric structure (Barnes et al., 2016). To determine how the *R24C* mutation affects UNC-27 function, we compared *unc-27(R24C)* mutants with a null allele generated by CRISPR/Cas9-mediated deletion of the entire *unc-27* coding sequence (referred to as *unc-27(0)*; see Methods). Consistent with the established role of UNC-27, *unc-27(0)* animals exhibited a marked reduction in body bend frequency (−66%, adjusted *p* <0.0001; **Fig. 2A**) accompanied by severe myofilament disorganization (**Fig. 2B**), highlighting the essential role of UNC-27 in maintaining sarcomere integrity. Similar to *unc-27(0)*, *unc-27(R24C)* mutants displayed a significant locomotor defect, with body bend frequency reduced by 50% relative to wild type (adjusted *p* <0.0001; **Fig. 2A**). This impairment was present in both young and aged animals. In contrast to *unc-27(0)* mutants, however, the sarcomeric organization of *unc-27(R24C)* animals was indistinguishable from that of wild type (**Fig. 2B**).

**Fig. 2.**
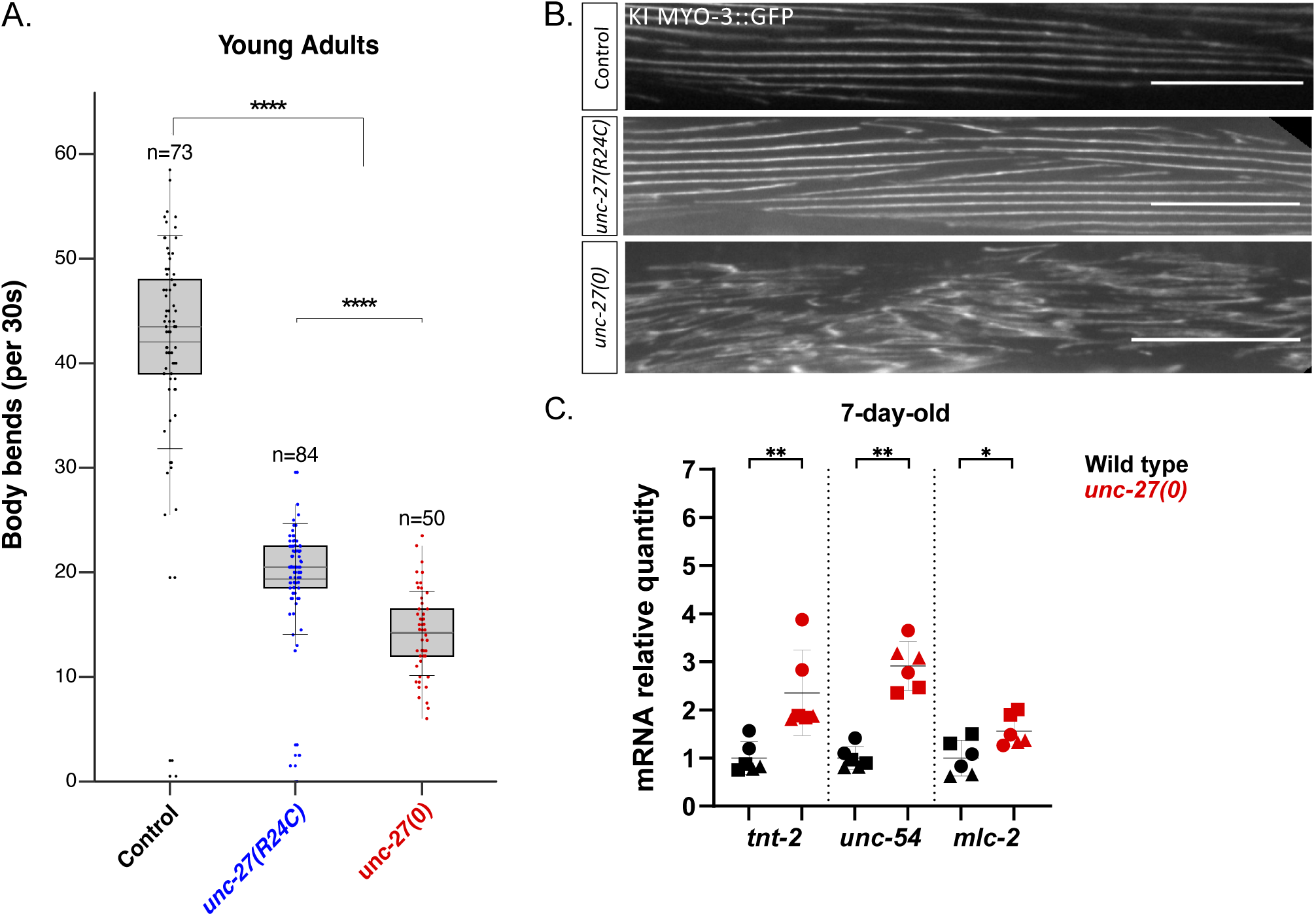
*unc-27(R24C)* only partially recapitulates the *unc-27* full loss-of-function phenotype. (A) Analysis of body bend frequency in young adults control, *unc-27(R24C)*, and null mutants. The number of animals per group is indicated above the bars (pooled from two experiments). Boxes represent the interquartile range (IQR); gray lines indicate medians; black lines indicate mean; whiskers represent SD. Statistical significance was assessed by Kruskal-Wallis with Dunn’s post hoc test (FDR adjusted); ****, *p*< 0.0001. (B) Representative images of MYO-3::GFP CRISPR knock-in expression in 2-day-old control, in *unc-27(R24C)* and *unc-27* null mutants. Scale bar, 25 μm. (C) Relative mRNA levels of *tnt-2*, *unc-54*, and *mlc-2* in 7-day-old adults *unc-27(0)* null mutants compared to wild-type animals. Each dot represents an individual replicate; biological and technical replicates are distinguished by different and identical shapes, respectively. Bars show median ± 95% confidence interval. Statistical significance was assessed by Mann-Whitney test; ****, *p*< 0.0001; **, *p*< 0.01; *, *p*< 0.05; ns, not significant.

Because *unc-27(R24C)* and *unc-27(0)* only partially share common phenotypes, we analyzed sarcomeric transcripts levels in *unc-27* null allele. Similar to *unc-27(R24C)*, complete loss of *unc-27* caused significant up-regulation of sarcomeric transcripts (**Fig. 2C**). Together, these findings demonstrate that UNC-27 has genetically separable roles in maintaining sarcomere structure and controlling sarcomeric gene expression.

### Sarcomeric disorganization but not altered muscle contraction is able to increase sarcomeric transcripts

Both *unc-27(R24C)* and *unc-27(0)* mutants exhibit reduced motility and increased sarcomeric transcripts. Since gene expression can be regulated by electrical activity and muscle contraction in vertebrate muscle (Schiaffino et al., 2007), we tested whether altered muscle activity is able to cause the up-regulation of sarcomeric transcripts. For this purpose, we used potassium channel mutants that affect muscle excitability. Potassium channels are key transmembrane proteins that regulate K⁺ flux, membrane repolarization and resting membrane potential. We chose the *unc-58(bln205)* and *twk-28(bln48)* mutants that display hypo- and hyper contracted muscle phenotypes, respectively (Andrini et al., 2024; Peysson et al., 2024).

While *twk-28(bln484)* mutants exhibit motility defects comparable to that of *unc-27(R24C), unc-58(bln205)* mutants are severely paralyzed (**Fig S1A, B**). RT-qPCR analysis revealed that sarcomeric transcript levels remained unchanged in *twk-28(bln484)*, but *unc-58(bln205)* mutants showed a modest increase in expression (**Fig. 3**). These finding initially suggested that the severe loss of motility in *unc-58* animals might contribute to the up-regulation of sarcomeric transcripts. However, further analysis revealed marked disorganization of both thin and thick filaments in *unc-58* mutants, a defect not observed in R24C mutants (**Fig S1C**). This observation led us to hypothesize that acute sarcomere disorganization, rather than altered contractility alone, triggers the transcriptional induction of sarcomeric genes in *unc-58(bln205)* mutants. To further explore this hypothesis, we examined additional mutants exhibiting varying degrees of thin- and thick-filament disorganization, including *unc-5*2, *unc-89,* and *unc-95* mutants (**Fig. S1B**). All mutants displayed elevated expression of *tnt-2, mlc-2,* and *unc-54,* although the magnitude of induction varied among strains (**Fig. 3**). Therefore, the disruption of sarcomere architecture appears sufficient to induce sarcomeric gene expression in multiple sarcomeric mutants, including *unc-27(0)* animals, while the loss of mobility alone is not. Thus, the transcriptional phenotype observed in R24C mutants is unlikely to arise from impaired muscle contraction.

**Fig. 3.**
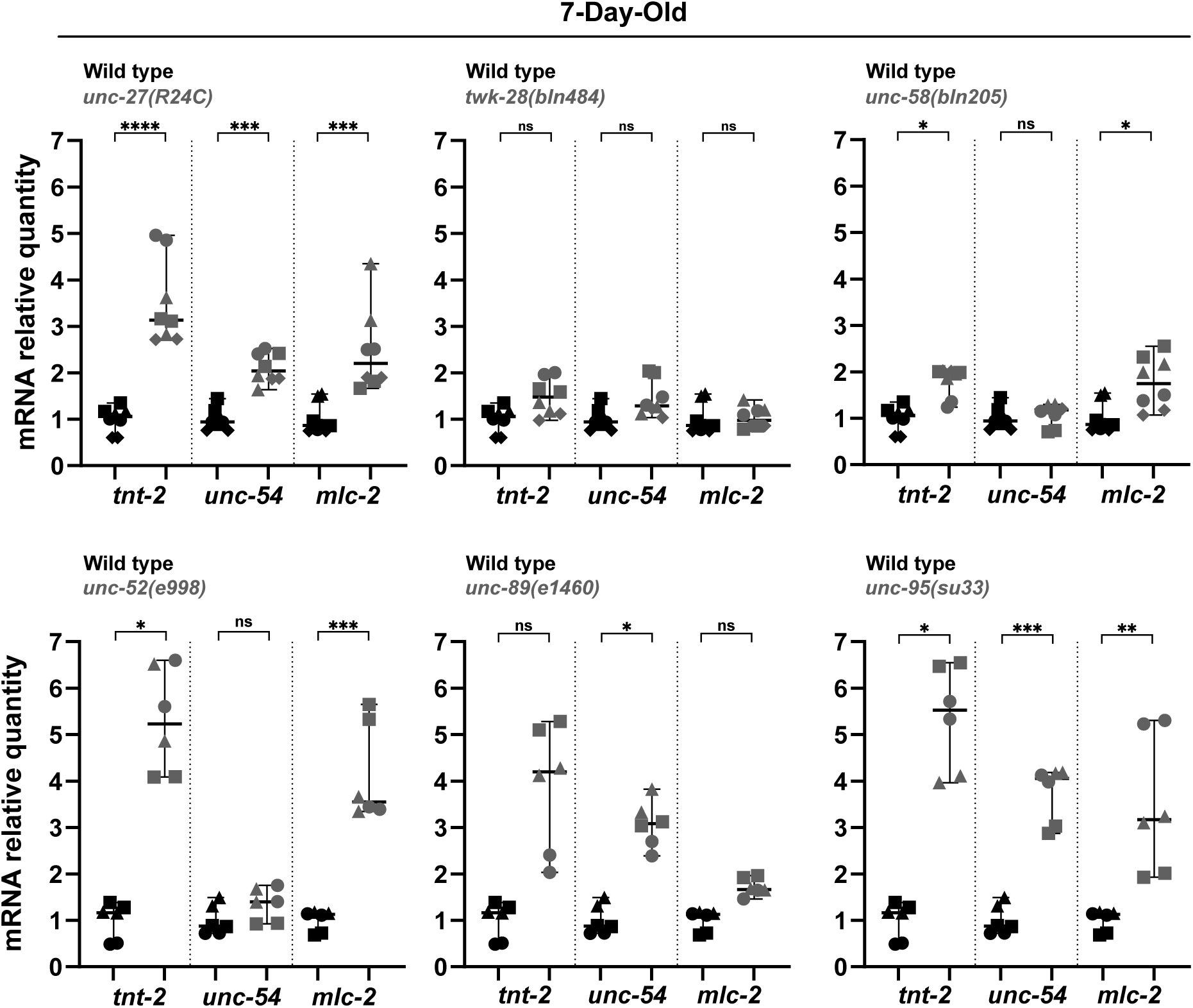
Sarcomere disorganization is associated to increased sarcomeric transcript levels. Relative mRNA quantities of *tnt-2*, *unc-54*, and *mlc-2* in 7-day-old adults for uncoordinated mutants associated with either potassium channels (*twk-28* and *unc-58*) or sarcomeric compartments (*unc-52*, *unc-89* and *unc-95*). Each dot represents an individual replicate; biological and technical replicates are distinguished by different and identical shapes, respectively. Bars show median ± 95% confidence interval. Statistical significance was determined by Kruskal-Wallis tests; ****, *p*<0.0001; ***, *p*<0.001; **, *p*<0.01; *, *p*<0.05; ns: not significant. SD, standard deviation.

### UNC-27 accumulates in muscle nuclei from early adulthood onwards

The *unc-27(R24C)* mutation results in a change from a conserved arginine to a cysteine at position 24 within the N-terminal region of the protein. Structural studies of the troponin complex reveal that the N-terminal region (amino acid 1 to 60) of troponin I lacks a defined secondary or tertiary structure, suggesting it is intrinsically disordered (Takeda et al., 2003; Yamada et al., 2020). However, several Nuclear Localization Signals (NLS) are predicted in this region (https://www.genscript.com/tools/psort) (NLS#1–3; **Fig. 4A**). Notably, the closest UNC-27 mammalian homolog, TNNI3, also harbors multiple predicted NLS in its N-terminal part (amino acids 11–50), along with an additional NLS in the C-terminal region (**Fig. 4A**). NLS sequences are typically required for active nuclear transport *via* importins of proteins larger than ∼40 kDa, whereas smaller proteins, such as UNC-27 (27 kDa), can passively diffuse through nuclear pores. To determine whether the predicted N-terminal NLSs of UNC-27 were functional, we generated a transgene encoding the first 60 amino acids of UNC-27 fused to a wScarlet::LacZ reporter, producing a ∼140 kDa fusion protein (**Fig. 4B**). When expressed as a multicopy array in muscle cells, the fusion protein localized to the muscle nuclei, indicating that the N-terminal domain of UNC-27 is sufficient to ensure active nuclear transport and that the predicted NLSs are functional (**Fig. 4C-D**).

**Fig. 4.**
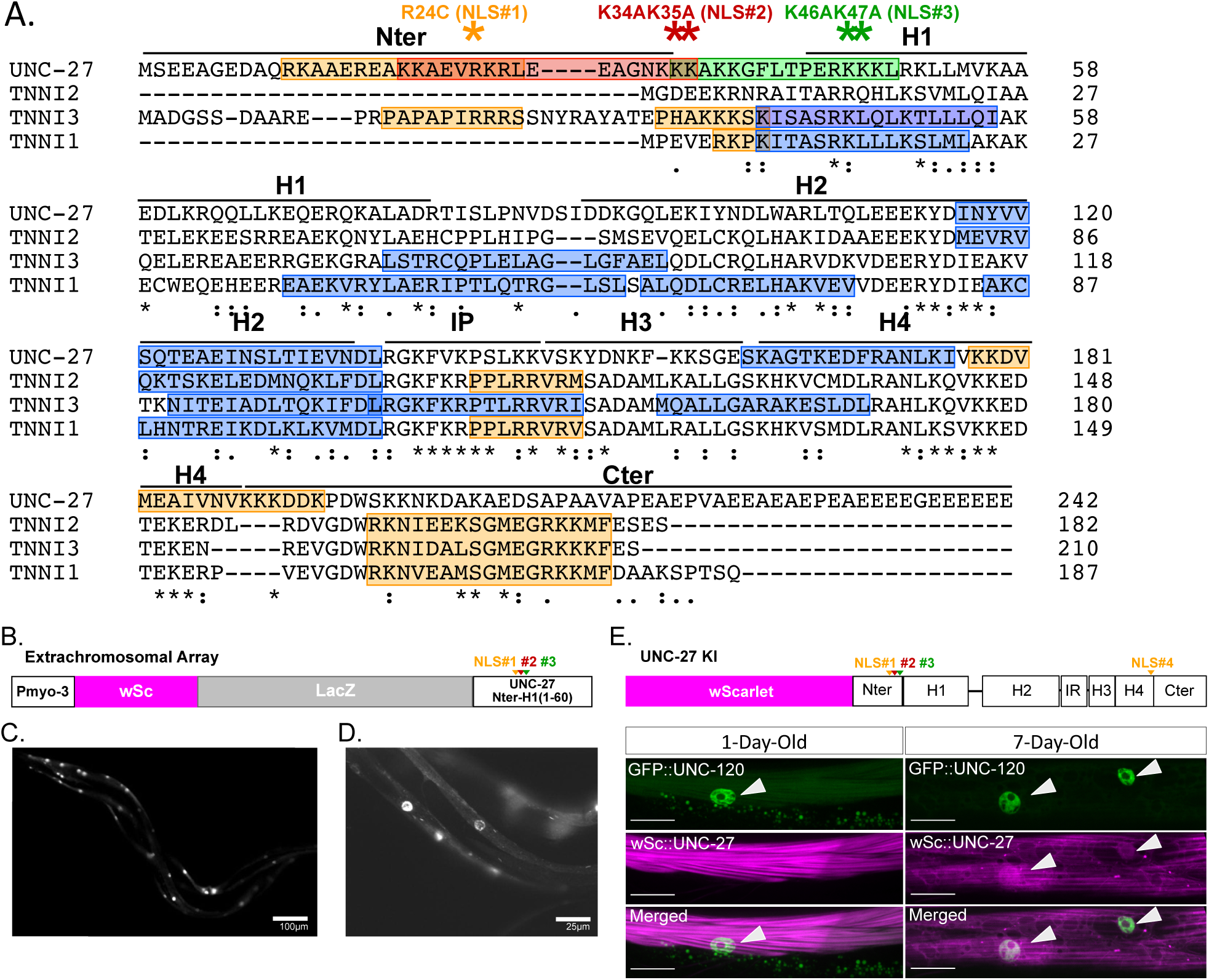
UNC-27/TNNI3 contains functional NLS motifs and accumulates in muscle nuclei during aging. (A) Clustal Omega sequence alignment of *C. elegans* UNC-27 and human skeletal (TNNI1-2) and cardiac (TNNI3) Troponin I isoforms. Yellow, green and red rectangles show predicted nuclear localization signals (NLS) NLS#1, #2 and #3 of UNC-27 respectively. Yellow and blue rectangles show predicted NLS and nuclear export signals (NES) respectively. Asterisks mark the CRISPR-induced mutation of NLS. Annotations show N-ter/C-ter domains, helices (H1-4), and the inhibitory peptide (IP). (B) Schematic of the 140 kDa chimeric wScarlet::LacZ::UNC-27 (Nter-H1) protein under the *myo-3* promoter. (C, D) Representative images of 1-day-old adult worms carrying the chimeric protein. (E) (Top) Schematic of wScarlet::UNC-27 knock-in. Arrows indicate predicted NLS positions. (Bottom) Representative images of age-dependent accumulation of wScarlet::UNC-27 in BWM nuclei at day 1 (left) and day 7 (right). White arrows indicate GFP::UNC-120/SRF positive nuclei, scale bars = 15 µm.

To assess the subcellular localization of UNC-27 under physiological expression levels, we inserted the fluorescent tag wScarlet at the 5’ end of the endogenous *unc-27* coding sequence. The tag did not disrupt animal motility or sarcomeric genes expression (**Fig. S2**). wScarlet::UNC-27 localized to the thin filaments of sarcomeres in one-day-old animals, as expected from its canonical role in the troponin complex (**Fig. 4E**). No nuclear signal of wScarlet::UNC-27 was observed at that age. However, by day 7 of adulthood, wScarlet::UNC-27 was also detected within muscle nuclei (**Fig. 4E**).

### UNC-27 nuclear localization is delayed in R24C mutant and suppressed in ΔNLS#1-3 mutant

NLSs are typically enriched in positively charged residues and we hypothesized that the R24C mutation, which results in the loss of a positive charge, impairs UNC-27 NLS#1 function. We compared the kinetics of UNC-27 nuclear localization between wild-type and R24C mutants with age. In wild-type animals, nuclear UNC-27 was first detected in a few nuclei on day 1 of adulthood, and the proportion of body wall muscle (BWM) nuclei exhibiting nuclear staining progressively increased, reaching a plateau by day 4 (**Fig. 5A**). In contrast, nuclear UNC-27(R24C) was detected only from day 3 of adulthood onward. (**Fig. 5A**). These data indicate that NLS#1 contributes to nuclear translocation of UNC-27. To assess the contribution of other NLSs, we generated a triple NLS mutant (ΔNLS#1-3: R24C, K34A, K35A, K46A, K47A, **Fig. 4A**). On day 4 of adulthood, no nuclear signal was detected in the ΔNLS#1-3 mutant, compared to ∼80% and ∼50% of stained BWM nuclei in WT and R24C mutant respectively (**Fig. 5B-C**). Together, these results demonstrate that age-dependent nuclear accumulation of UNC-27 is a regulated process involving multiple NLS motifs.

**Fig. 5.**
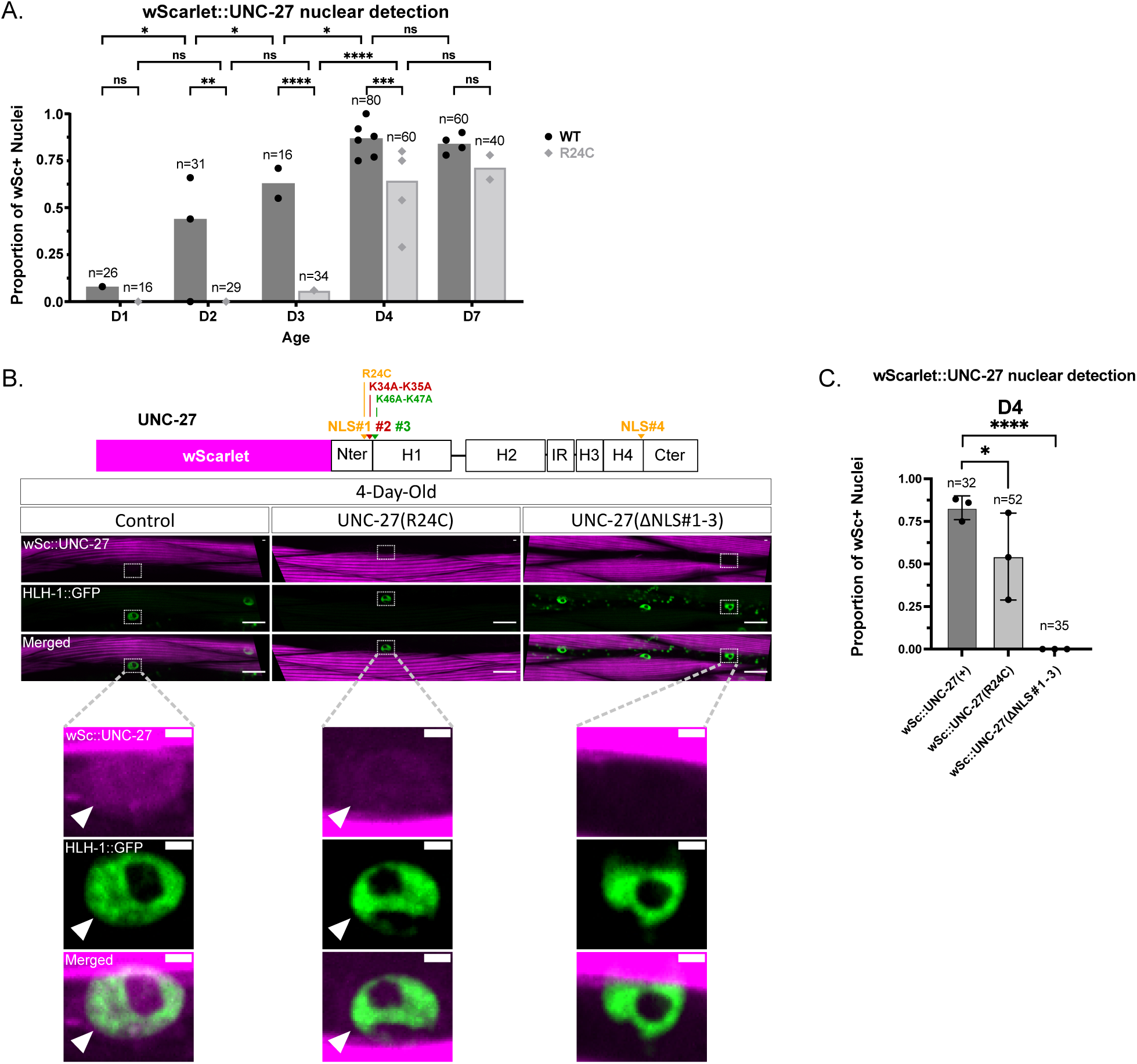
Mutations in the NLS prevent age-associated nuclear accumulation of UNC-27. (A) Quantification of the percentage of BWM nuclei positive for wSc::UNC-27 from day 1 to day 7 in WT and R24C mutants. Dots represent means from at least 5 worms per biological replicates; n denotes total nuclei screened. Statistical significance was determined by two-sided Fisher’s exact test; *\*\*\*\*, p<*0.0001; *\*\*\*, p<*0.001; *\*\*, p<*0.01; *, *p*<0.05; ns, not significant. (B) Schematic of wScarlet::UNC-27 knock-in, arrows indicate predicted NLS positions. Confocal images of wSc::UNC-27 in 4-day-old control, R24C, and ΔNLS#1-3 mutants. Scale bars: 15 μm (top) and 2 μm (bottom). BWM nuclei are detected with HLH-1::GFP KI. In the bottom panels, arrows indicate positive nuclei. (C) Quantification of nuclear detection in day-4 adults. Dots represent means from at least 5 worms per biological replicates; n denotes total nuclei screened. Statistical significance was determined by two-sided Fisher’s exact test; *\*\*\*\*, p<*0.0001; *, *p*<0.05; ns, not significant.

### Nuclear UNC-27 downregulates sarcomeric transcripts

Our data showed that the regulation of sarcomere transcripts correlates with UNC-27 nuclear accumulation and suggest that nuclear UNC-27 may act as a repressor of sarcomere gene expression. We first asked whether complete removal of UNC-27 from muscle nuclei may further enhance up-regulation of sarcomere gene expression. Mutating all three NLS regions is indeed associated with a further increase in *tnt-2* transcriptional reporter expression compared to mutation of NLS#1 alone (R24C mutants, **Fig. 6A-B**), showing that UNC-27’s impact on *tnt-2* transcripts is modulated by its nuclear translocation.

**Fig. 6.**
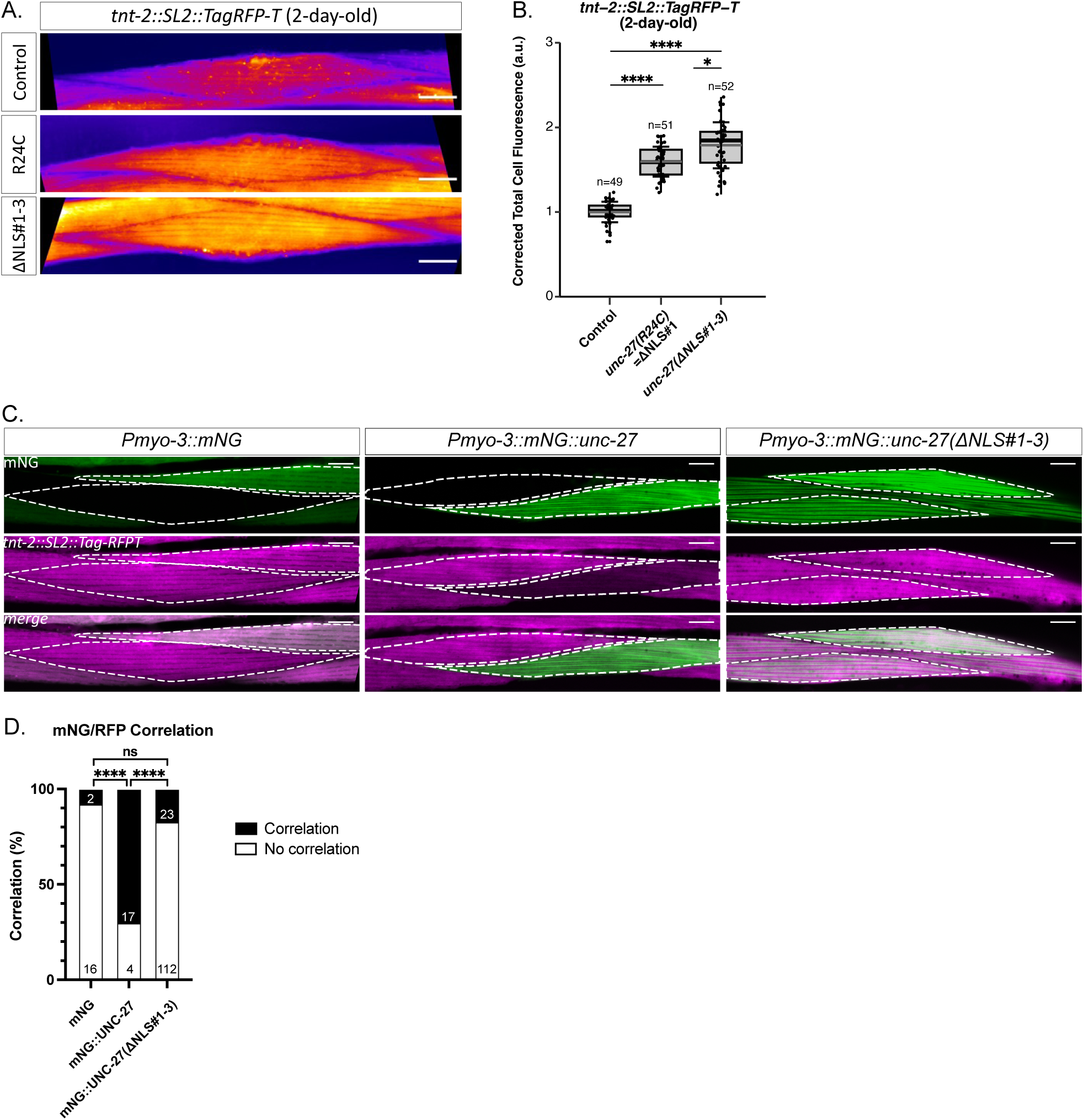
Nuclear UNC-27 cell-autonomously represses *tnt-2* expression. (A) Representative images of the *tnt-2* transcriptional fluorescent reporter in control, R24C and ΔNLS#1-3 worms. (B) Quantification of reporter fluorescence, where each dot represents an individual cell. Boxes indicate interquartile range; center lines indicate medians; whiskers indicate SD. Statistical significance was determined by Kruskal-Wallis with Dunn’s test; ****, *p* <0.0001; *, *p*<0.05; ns, not significant. (C, D) Mosaic analysis of muscle-specific UNC-27 over-expression. (C) Representative images of 2-day-old mosaic animals expressing muscle-specific mNG, mNG::UNC-27 WT, or mNG::UNC-27(ΔNLS#1-3). Dashed outlines indicate adjacent cells scored first for differential mNeonGreen expression and subsequently for TagRFP-T intensity. (D) Quantification of the mNG/RFP correlation in mosaic worms. For the mNG::UNC-27(ΔNLS#1-3) overexpression, observations of 5 independent strains have been pooled. The number of pair of cells scored are indicated inside the bars. Statistical significance was determined by two-sided Fisher’s exact test; ****, *p*<0.0001; *, *p*<0.05; ns, not significant.

To determine whether UNC-27 overexpression in a WT background is sufficient to reduce *tnt-2* transcript levels, we generated transgenic mosaic animals expressing wild-type mNeonGreen-fused (mNG) UNC-27 protein (WT) in a *tnt-2::SL2::tagRFP-T* knock-in transcriptional reporter background (**Fig. 6C**). These animals carry extrachromosomal multicopy transgenes that are randomly lost during cell divisions, resulting in variability in transgene doses between cells of the same individual. We then examined the *tnt-2* reporter-associated fluorescence in cells expressing different levels of mNG::UNC-27 and compared it to that in transgenic worms expressing the fluorescent protein mNG alone. At day 2 of adulthood, 70% of BWM cells overexpressing wild-type mNG::UNC-27 exhibited reduced *tnt-2* reporter fluorescence, compared to cells with low level of mNG::UNC-27 (**Fig. 6C-D**). In contrast, no decrease in *tnt-2* reporter expression was observed in cells expressing high level of mNG alone (**Fig. 6C-D**). Furthermore, transgenic animals expressing mNG::UNC-27(ΔNLS#1-3) behaved like mNG controls (**Fig. 6C-D**). Together, these findings showed that UNC-27 acts cell-autonomously as a repressor of sarcomeric gene expression and indicate that its nuclear localization is required for the downregulation of sarcomeric transcripts.

### Transcriptomic landscape of UNC-27 ΔNLS#1-3 mutant

We then investigated whether the nuclear UNC-27 protein influences transcripts other than sarcomeric genes during aging. To this end, we analyzed the transcriptome of wild-type worms and NLS mutants (UNC-27(ΔNLS#1-3)), at the pre-adult L4 stage, young-adult stage and in 7-day-old adult animals across four biological replicates. As an initial step, we estimated the biological age of each sample from its transcriptomic profile using RAPToR, which infers developmental age by comparing gene expression patterns to a reference developmental time series (Bulteau & Francesconi, 2022). This approach enables more precise isolation of mutation-specific effects by eliminating transcriptomic variations linked to subtle developmental differences between experimental replicates (Bulteau & Francesconi, 2022). Our analysis revealed that on day 7 of adulthood, the mutants were predicted to be younger than wild-type animals, while their estimated ages were comparable at earlier stages (L4 and young adult stages) (**Fig.7A**). Analysis of differentially expressed genes (DEG; fold change > 2, p-value < 0.01) identified 83 up-regulated and 69 down-regulated genes in 7-day-old mutants compared to wild-type animals (**Fig. 7B**). Muscle-expressed genes account for over 70% (59/83, **Table S1**, CenGen Consortium) of the up-regulated genes. In contrast, approximately 70% of down-regulated genes (48/69, **Table S1**) are expressed outside muscle tissue, while only 11% are muscle-enriched (**Table S1**). These findings suggest that nuclear UNC-27 primarily acts as a transcriptional repressor in muscle.

**Fig. 7.**
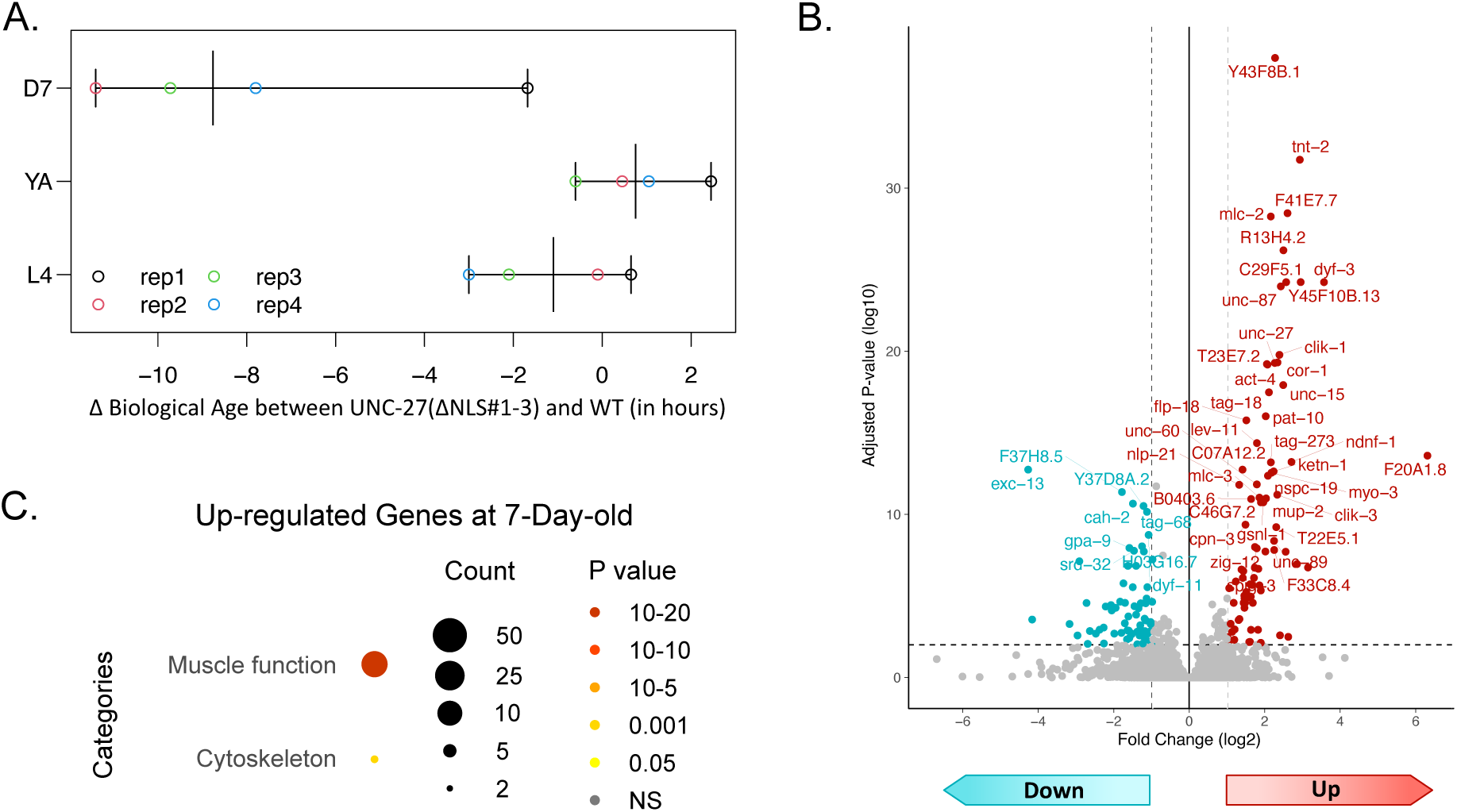
Aged UNC-27 ΔNLS mutants exhibit a reduced biological age associated with selective remodelling of the muscle transcriptome. (A) Difference in estimated biological age, as determined by RAPToR, between wild-type (WT) and ΔNLS#1–3 mutants at the L4 larval stage, young adulthood, and day 7 of adulthood. (B) Volcano plot showing differentially expressed genes (DEGs) between day-7 ΔNLS#1–3 mutants and WT animals. Differential expression thresholds were set at log₂ Fold Change (FC) > 1 and *p_adj_* < 0.01 (indicated by dotted lines). (C) WormCat enrichment analysis of genes upregulated in day-7 ΔNLS#1–3 mutants, grouped by Category 1 annotations. Statistical significance was assessed using a Fisher’s exact test with Bonferroni correction *p*< 0.05. Abbreviations: FC, fold change; WT, wild type. For further details on the DEG, please refer to Table S1.

Gene set enrichment analysis using WormCat (Holdorf et al., 2020) revealed no enrichment among down-regulated genes. In contrast, up-regulated genes were significantly enriched for muscle function (16 out of 36 genes with assigned functions, **Fig. 7C**), including myosin heavy and light chains, troponin complex subunits, tropomyosin, alpha-actinin, and zyxin, as well as for cytoskeleton-related genes (9 out of 36).

Overall, these data support the age-dependent role of nuclear UNC-27 and indicate that it primarily acts as a repressor of sarcomeric genes in muscle. Additionally, preventing its nuclear translocation appears to trigger secondary signaling, as evidenced by the regulation of genes expressed outside muscle tissue (with 25% and 63% of neuronal enriched genes among up-regulated and down-regulated genes respectively, **Table S1**).

### Preventing the nuclear translocation of UNC-27 improves protein homeostasis and protects against age-related mitochondrial fragmentation

A stereotyped sequence of conserved subcellular changes (i.e. biomarkers) defines muscle aging in *C. elegans* (Mergoud Dit Lamarche et al., 2018), however, the causal relationships between these different biomarkers remain poorly understood. We thus sought to determine whether other age-related changes might be affected by the R24C and ΔNLS#1-3 mutations. We first assessed muscle proteostasis using a strain expressing an aggregate-prone YFP fused to 35 polyglutamine (polyQ) repeats, under the control of a muscle-specific promoter. This strain, also used as a model to mimic Huntington’s disease, is commonly employed to evaluate muscle cells proteostatic capacity. Young animals exhibited a diffuse YFP fluorescence pattern, whereas two-day-old adults displayed visible aggregates, the number of which increased with age (**Fig. 8A-B**). The R24C mutant showed fewer aggregates up to day 7 of adulthood. Interestingly, in the ΔNLS#1-3 mutant, the formation of aggregates is even further delayed (**Fig. 8A-B**). Thus, inhibiting the nuclear localization of UNC-27 improves protein homeostasis during early adulthood.

**Fig. 8.**
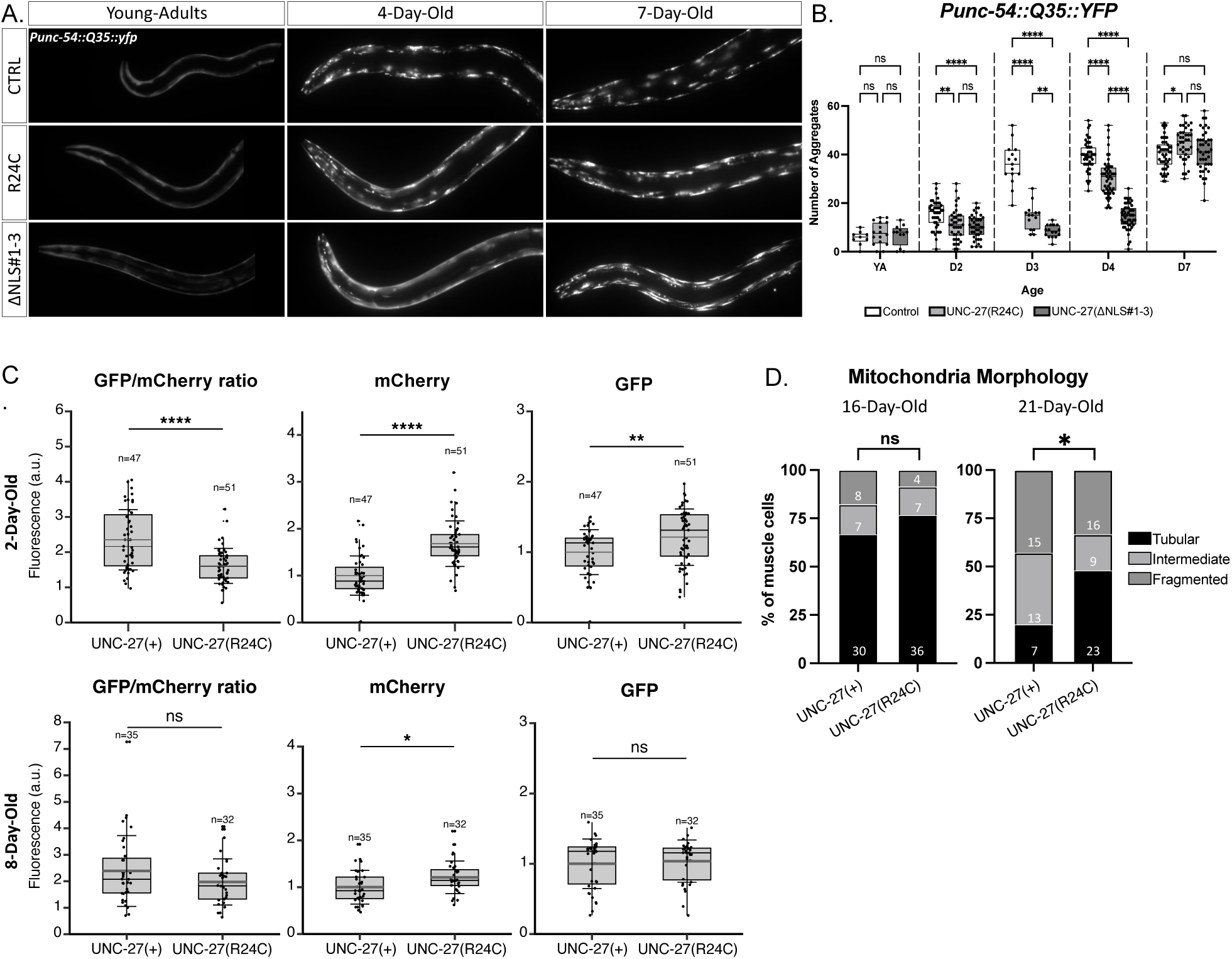
Preventing UNC-27 nuclear translocation improved protein homeostasis and delayed mitochondria fragmentation. (A, B) Analysis of Q35::YFP protein aggregates formation in WT, R24C, and ΔNLS#1-3 animals. (A) Representative images of Q35::YFP aggregation in BWM in young adults, 4-day-old or 7-day-old adults. (B) Quantification of aggregates number in the anterior BWM of the worms. Each dot represents individual worms. Boxes indicate IQR; black lines indicate medians; whiskers indicate min/max values Statistical significance was determined by Tukey’s test. (C) Assessment of autophagic flux via quantification of muscle-specific SQST-1::GFP::mCherry levels and GFP/mCherry ratio in WT, R24C, and ΔNLS#1-3 animals at day 2 and day 8 of adulthood. Each dot represents individual worms. Boxes indicate IQR; center lines indicate medians (gray) or means (black); whiskers indicate SD. Statistical significance was determined by two-sided Wilcoxon-Mann-Whitney. (D) Analysis of mitochondrial morphology using a muscle-specific reporter (TOMM-20N::Scarlet) in WT and R24C animals at day 16 and day 21 of adulthood. Inside the bars, n represents the number of animal per categories. Statistical significance was determined by Chi-squared tests; *\*\*\*\*, p<*0.0001; *\*\*\*, p<*0.001; *\*\*, p<*0.01; *\*, p<*0.05; ns, not significant.

We then asked whether this delay may be linked to an improvement of autophagic activity. We and others have previously shown that age-related blockade of autophagy can be detected in BWM as soon as 7 days of adulthood (Chang et al., 2017; Mergoud Dit Lamarche et al., 2018). We used the dual fluorescent autophagy reporter SQST-1/p62::mCherry::GFP to monitor autophagic flux *in vivo* (Villalobos et al., 2023). At day 2 of adulthood, R24C mutant displayed reduced GFP/mCherry ratio compared to control worms, indicating an increase of autophagic flux (**Fig. 8C**). However, at day 8 of adulthood, GFP/mCherry ratio of R24C mutant was similar to control. Overall, these findings suggest that preventing the nuclear translocation of UNC-27 improves protein homeostasis through the direct or indirect regulation of autophagic flux in early adulthood, but does not delay the onset of autophagy blockade.

Another key biomarker of muscle aging is progressive mitochondrial fragmentation. To assess this phenotype, we used a single-copy mitochondrial reporter consisting of the N-terminal domain of TOMM-20 fused to wScarlet (Roy et al., 2022). Unlike overexpressed reporters that induce fragmentation artifacts (Fabrizio et al., 2024), this reporter reveals the late-onset mitochondrial fragmentation that occurs during normal aging. Using this system, we found that mitochondrial fragmentation was delayed in R24C mutants compared to wild-type animals. Notably, 21-day-old R24C mutants exhibited significantly less fragmentation than age-matched controls (**Fig. 8D**). Together, these results indicate that preventing UNC-27 nuclear translocation preserves sarcomeric transcript levels and delays multiple hallmarks of muscle aging, including proteostasis decline and mitochondrial fragmentation.

## Discussion

In this study, we conducted the first visual genetic screen aiming at identifying regulators of muscle aging. This strategy led to the discovery of a new *unc-27* mutant allele. Its characterization uncovered an unexpected nuclear role for a sarcomeric protein in regulating sarcomeric gene expression in aged animals, with an impact on several subcellular hallmarks of muscle aging.

### A novel nuclear function of UNC-27 in the regulation of muscle aging biomarkers

We demonstrated that UNC-27 is actively transported into the nuclei of muscle cells with age. Disrupting this transport delayed the age-related decline of sarcomeric transcripts, improved early protein homeostasis, and postponed mitochondrial fragmentation. While transcriptomic analyses support a role for nuclear UNC-27 in repressing sarcomeric gene expression with age, they did not identify downstream targets directly linked to protein or mitochondrial homeostasis, suggesting that these phenotypes may result from the up-regulation of sarcomeric genes. Sarcomeric proteins are the most abundant proteins in muscle cells, and their increased levels impose a burden on the proteostasis network (Dorsch et al., 2019). Surprisingly, we observe an improvement in protein homeostasis at the onset of adulthood. This suggests that elevated sarcomeric protein levels may trigger an adaptive response, enhancing proteostasis capacity during the critical developmental stage of peak egg production. However, this early compensatory mechanism, which we show depends at least in part on increased autophagic flux, is transient. It ultimately fails to prevent the subsequent blockade of autophagy, which emerges during the first week of adulthood.

The maintenance of sarcomeric proteins may also influence mitochondria homeostasis. Recent work in *Drosophila melanogaster* has demonstrated that sarcomeric proteins and mitochondrial morphogenesis are coordinated through a mechanical feedback loop where sarcomere assembly drives mitochondrial intercalation and elongation between myofibrils (Avellaneda et al., 2021). Supporting this model in *C. elegans*, we observed that *unc-27* null mutants, which exhibit sarcomere disorganization, also display constitutive mitochondrial fragmentation (data not shown). These findings suggest a previously underappreciated role for sarcomeric proteins in maintaining mitochondrial homeostasis during aging in *C. elegans*.

Finally, we showed that preventing UNC-27 nuclear translocation triggers cell non-autonomous signaling, as evidenced by the increased expression of genes associated with cell types other than muscle in our transcriptomic analysis, including neuronal genes. Taken together, these findings highlight the contribution of UNC-27 to both early and late changes associated with muscle aging, acting through cell-autonomous mechanisms as well as through systemic, non-cell-autonomous pathways.

### Conservation of the nuclear role of UNC-27/TNNI3 in the regulation of sarcomeric genes in mammals

Three isoforms of troponin I have been identified in mammals (TNNI1 to 3). The closest orthologue of *unc-27* is the cardiac isoform, TNNI3. Similarly to UNC-27, it contains an N-terminal domain with an NLS (Fig. 4A and Barnes et al., 2016), which is absent from the other two troponin I isoforms expressed in skeletal muscle (TNNI1 and TNNI2). Bergmann and colleagues were the first to report the detection of a nuclear pool of TNNI3 in cardiomyocytes isolated from the human heart (Bergmann et al., 2009, 2011). Nuclear localization of TNNI3 was later observed in cardiomyocytes derived from rat mesenchymal cells (Asumda & Chase, 2012). Furthermore detection of nuclear troponin I has also been reported in non-muscle cells, as illustrated in the early stages of *Drosophila melanogaster* embryonic development, as well as in S2 insect cells (Sahota et al., 2009). However, the impact of nuclear TNNI3 on the transcriptome remains systematically unexplored. Using a candidate approach, Tian’s group proposed that *Atp2a2* and *PDE4D* might be regulated by TNNI3 in mouse cardiomyocytes (Lu et al., 2022; Zhao et al., 2020). Although the *C. elegans* orthologs of these genes (*sca-1* and *pde-4*) are conserved and expressed in muscle, their expression was unaffected by the loss of nuclear UNC-27 (Supplementary Table 2). Whether TNNI3 also regulates sarcomeric genes remains an open question.

Overall, our results provide novel insights into the biological significance of nuclear troponin and lay the groundwork for future investigations into its potential roles in aging and muscle-related pathologies.

### A signaling role for the sarcomeric proteins in coordinating myofibrillar organization with transcriptional regulation

We showed that the subcellular localization of UNC-27 is a dynamic process involving several NLS for its nuclear transport. Notably, the protein also contains a predicted nuclear export domain (NES), suggesting nucleocytoplasmic shuttling between the nuclear and cytoplasmic/sarcomeric pools of UNC-27. What triggers nuclear accumulation with age remains an open question. A number of post-translational modifications of TNNI3 have been reported, including SUMOylation, phosphorylation and proteolytic cleavage of the N-terminal domain, yet their impact on the nuclear transport of TNNI3 is unknown (Gao et al., 1997; Janssens et al., 2018; Sahota et al., 2009). Cleavage of UNC-27 does not appear to be involved in the regulation of its nuclear function, as UNC-27 fused to wScarlet can be detected in nuclei with age regardless of whether it is tagged at the N- or C-terminus (**Fig.4-5** and data not shown). However, the UNC-27 protein contains multiple sequences that are predicted to be the target of several post-translational modifications and might regulate its nuclear localization. This will require further investigations.

Our data assign a central role to UNC-27 in the repression of several sarcomeric genes that are regulated as a module during aging, although the molecular mechanisms involved remain unknown. The absence of predicted DNA or RNA binding site in the protein suggests that it may regulate gene expression indirectly, for example by serving as a scaffold for chromatin proteins or transcription factors. Overexpression experiments indicate that UNC-27 is sufficient to regulate these genes cell-autonomously, suggesting that its partners are not limiting. Interestingly, a previous study described the nuclear accumulation of mutated troponin T (TNT) in cardiomyocytes derived from iPS cells of a patient with cardiomyopathy. TNT was shown to interact with conserved histone demethylases (Wu et al., 2015). Two TNTs, TNT-2 and MUP-2 are expressed in *C. elegans* muscle. Since troponin I and troponin T are part of the same complex in sarcomeres, whether UNC-27 also functions with these proteins in the regulation of sarcomeric genes is an open hypothesis.

Our data show that signaling pathways link sarcomere integrity and sarcomeric genes expression. Severe disorganization of myofilaments in mutants prevents the repression of sarcomeric genes. Whether this is triggered by specific sarcomere integrity sensors or by non-physiological activation of stress responses has not been investigated. In the physiological context of aging, troponin I/UNC-27 plays a central role in downregulating a large set of muscle-specific genes predominantly involved in muscle contraction. Paradoxically, impairing this function not only delays the cellular hallmarks of muscle aging, but also partially slows transcriptional signatures of aging at the organismal level (Fig.7A). These findings could lead to the identification of new regulatory networks required to control gene expression at multiple levels. Strikingly, similar to the single amino acid substitution R24C within the conserved NLS1 sequence of UNC-27 that disrupts its nuclear function, human TNNI3 harbors the equivalent pathogenic R21C mutation, associated with hypertrophic cardiomyopathy (Fahed et al., 2020; Gomes et al., 2005). These findings raise the possibility that altered regulation of the nuclear pool of TNNI3 may contribute to disease pathogenesis in addition to its established role in contractile dysfunction.

## Acknowledgement

This work was supported by the French National Research Agency (ANR-21-CE14-0026-01) and AFM-Téléthon (JCD-2024 Scientific Call). We thank Le Centre d’Imagerie Quantitative Lyon-Est (LyMIC-CIQLE, Lyon, France) imaging facility for support and access to equipment, and Camilla Luccardini for technical assistance. We thank ProfileXpert plateforme (IBiSA, SFR Santé Lyon Est UAR3453 CNRS/US7 INSERM), particularly Severine Croze, and Guillaume Marcy (LABEX Cortex and Université Lyon 1) for the RNA sequencing and analysis. We thank the Caenorhabditis Genetic Center (which is funded by NIH Office of Research Infrastructure Programs, P40 OD010440) for strains. We thank Dr. Jonathan Enriquez and Dr. Anna Mattout for their insights on the project.

## Declaration of generative AI and AI-assisted technologies in the writing process

During the preparation of this work the authors used ChatGPT - OpenAI and DeepL in order to improve the readability and language of the manuscript. After using these tools, the authors reviewed and edited the content as needed and take full responsibility for the content of the publication.

## Method

### 1. Strains and genetics

All C. elegans strains were grown at 20°C on nematode growth medium (NGM) agar plates with Escherichia coli OP50 as a food source. All strains were originally derived from the wild-type Bristol N2 strain. A complete list of strains used in this study is provided in Table Supp 2.

### 2. EMS screen

Worms were collected and washed with M9 buffer (3 g of KH2PO4, 6 g of Na2HPO4, 5 g of NaCl and 0.25 g of MgSO4·7 H2O, distilled water up to 1 L) from 10 plates containing a majority of L4 (4^th^ larval stage). Mutagenesis was performed by incubating 4 mL of M9 containing the pellet of worms (P0) with 20 μL of EMS (SIGMA M-0880) for 4 h with agitation. Worms were then washed 5 times in 15 mL of M9 and put back on NGM plates with OP50. F1 progeny was cloned on new fresh plates for 2 days at 20°C, and transferred again at 15°C. From the 20°C plate, 10 to 20 F2 per F1 at the L4 stage were isolated in a well of a 24 well-plate containing 10µM of 5-FU and transferred at 25°C. Five days later, the 20 F2 were screened for worms with bright signal from the BWM, under a Nikon AZ100 multizoom microscope. When a mutant with high fluorescence was identified in the well, the 15°C F1 plate was screened for homozygous mutants.

### Mapping, whole genome sequencing and validation of mutations

Homozygous mutants were backcrossed twice to the parental strain to remove non-causal mutations. After backcrossing, genomic DNA was extracted for whole-genome sequencing (Eurofins and Novogene kit), from the pool of 5 independent F2 from the outcross for each strain. Identification of causative EMS-induced mutations was performed using in-house built-in scripts. Candidate mutations were validated by recreating the mutation in the parental background with CRISPR/Cas9.

### 3. CRISPR/Cas9 genome engineering

All knock-in alleles were generated according to Ghanta and Mello 2020. crRNA was designed with the Benchling software and synthesized by Integrated DNA Technologies (IDT). A mix containing 2.8 μL crRNA (34 μM), 5 μL tracrRNA (18 μM, IDT #1073190) and 0.5 μL Cas9 nuclease (10 mL/mg, IDT #1081058) was incubated for 15 min at 37°C. For the generation of deletions, 2 crRNAs were used, at both ends of the deleted sequence. In this case, the mix contained 1.4 μL of each crRNA. A mix of 2.2 μL single strand repair template (1 μg/μL) or 500 ng double strand repair template, with 800 ng of pRF4 plasmid and molecular biology grade water up to 20 µL was added to the mix. This mix was injected in adult worms. Worms with a roller phenotype in the next generation were isolated and tested by PCR. All gene editions were confirmed by sequencing. Finally, the edited strains were outcrossed at least once. A list of CRISPR alleles generated in this study and corresponding crRNAs is provided in Table Supp. 2.

### 4. Generation of mosaic animals

*tnt-2::SL2::TagRFP-T (FS410 strain)* animals were injected with a mix of 5 ng.µL^-1^ of plasmid (Table Supp 3), 20 ng.µL^-1^ of pRF4, complemented with 75 ng.µL^-1^ of 1kb+ (Invitrogen).

Animals carrying the extrachromosomal transgene were selected on the roller phenotype.

### 5. Plasmid construction

The plasmids constructed for this study are described in Table Supp. 2. All constructs were verified by Sanger sequencing (Eurofins).

### 6. Live Confocal Microscopy

Animals were placed on 20 µM 5-FU plates at the L4 larval stage, and grown for the indicated ages. For live imaging, synchronized adult hermaphrodites were mounted blindly on 2% agarose (w/v in water) dry pads immersed in 5% Polybead microspheres (#00876-15 Biovalley) diluted in M9 buffer. Confocal images were acquired under an Andor spinning disk system (Oxford Instruments) installed on a Nikon-IX86 microscope (Olympus) equipped with a 40X/NA1.3 oil immersion objective and an Evolve EMCCD camera.

For nuclear localization scoring, BWM nuclei were detected using either *fs442 gfp::unc-120* or *kagSi hlh-1::gfp* knock-in. When the cell orientation placed the nucleus outside the sarcomere, images were acquired as a stack of optical sections (0.5 µm apart). Scoring of wScarlet::UNC-27–positive nuclei was performed in a double-blind manner, with the experimenters unaware of both the strain identity and image source. Approximately 5 cells per animal were assayed, with at least 5 worms per replicate.

For quantification of *tnt-2* transcriptional reporter fluorescence, images of body wall muscle (BWM) cells along the entire body were acquired as Z-stacks, with optical sections spaced 0.5 μm apart. Signal intensity was quantified using a custom macro in Fiji (ImageJ). Briefly, Z-stacks were first cropped to isolate individual cells within rectangular regions encompassing each entire cell. These stacks were then projected along the Z-axis using sum projections. For each projected image, a region of interest (ROI) corresponding to the BWM cell was manually outlined, and the the sum of pixel intensities within the ROI) was measured (Raw Integrated Density). Background fluorescence corresponds to the mean background intensity from small ROIs placed outside muscle cells, within the control worms. The Corrected Total Cell Fluorescence (CTCF) was then calculated as: *CTCF = Raw Integrated Density – (mean background intensity in control × ROI area)*. This approach ensured consistent and standardized background correction across all samples.

### 7. Autophagy, proteostasis and mitochondrial network assessment in BWM

For *rmIs132*[*Punc-54::Q35::YFP*] aggregates quantification, and *krSi134*[*Pmyo-3::tomm-20 N::wScarlet*] mitochondrial network assessment, worms were synchronized since the L4 stage, using 5-FU 20 µM plates. Animals of indicated ages were blindly mounted as described above, and imaged under Axioscop compound microscope (Zeiss) equipped with Neofluar 10X/NA 0.30 air immersion and 63x/NA 1.25 oil-immersion objectives and a EMCC CoolSnap HQ (Photometrics) camera.

For mitochondrial network, images from the posterior BWM cells of each worm were acquired. Strains scoring and image analysis were performed blind. Cells with long interconnected mitochondrial networks were noted interconnected; cells with a combination of interconnected mitochondrial networks along with some smaller fragmented mitochondria were classified as interrupted; cells with sparse small round mitochondria were classified as fragmented.

For *rmIs132* [*Punc-54::Q35::YFP*] aggregates quantification, one focal plane of the anterior half of each worm was acquired. Counting of aggregates was performed in FIJI. Briefly, images were converted in binary 8-bit images, and aggregates were isolated with thresholding (Intermode algorithm) and particles analysis for counting.

For *louIs10* [*Pmyo-3::sqst-1::mCherry::GFP*] quantification, worms were imaged under Thunder Imager Live Cell inverted microscope (Leica) with a HC PL Fluotar 10X/NA 0.30 objective. Fluorescence quantification of each channel was performed automatically in FIJI. Briefly, for each image, individual worms were isolated and channels split. Each channel underwent background subtraction, median filtering, and thresholding to generate binary masks. Aggregates were detected using “Analyze Particles” and their integrated fluorescence were measured.

### 8. RNA extraction and Reverse transcription– qPCR

#### RNA extraction

Worms were synchronized by egg-laying (20 adults/plate for 1 hour), and exposed to 20 µM 5-FU from the L4 stage. 300 young-adults or 7-day-old worms were collected and washed twice in M9 buffer, before storage at -80°C in 1 mL TRIzol (Invitrogen). RNAs were extracted using phase separation followed by phenol−chloroform purification (QIAgen MaXtractTM High Density). Quantification of total RNA was done on a NanoDrop 1000 spectrophotometer (ThermoScientific) and their quality was assessed by RIN measurement (RNA integrity number) using a Agilent 2100 Bioanalyzer. DNase treatment was performed with the TurboDNA-free kit (Thermo Fisher Scientific).

#### RT-qPCR

cDNA was synthesized with the iScript cDNA synthesis kit (Bio-Rad) following provider’s instructions. qPCR reactions were performed using iTaq Universal SYBR (Biorad) in a CFX96 Realtime System C1000 Thermal Cycler (Bio-Rad). The qPCR protocol consisted of an initial step at 95°C for 4 min, followed by an amplification program during 40 cycles (30 sec at 95°C, 30 sec at 60°C and 30 sec at 72°C). Product amplification was verified with Melting-curve analyses. Normalization was done with the geometrical mean of three housekeeping genes (*tba-1*, *pmp-3* and Y45F10D.4). All primers were designed using NCBI Primer - BLAST and selected to generate amplicons with a length of 100−200 bp. Standard curves were generated for each primer set to calculate the efficiency of each set. Only primer sets with an efficiency of 1.8−2.1 were used for RT qPCR. RT qPCR experiments were repeated at least three times using independent biological samples with two technical duplicates. Primers used are reported in Table Supp. 2.

### 9. Whole worm RNA sequencing

#### RNA extraction

total RNA was extracted as described above.

#### Library preparation

Library preparation and RNA sequencing were performed at the ProfileXpert platform (UCBL, Lyon). Quality (RIN ≥ 8) of samples was checked by Fragment Analyzer (Agilent) and RNA was quantified by Nanodrop and by Quantifluor RNA kit (Promega). First, mRNA was enriched from 300 ng of total RNA, then library preparation was realized with the MGIEasy RNA Directional Library Prep Set (MGI). Quality of libraries were checked by Fragment Analyzer (Agilent) and quantified by Qubit 1X dsDNA HS Assay Kit. After circularization in ssDNA, DNA NanoBalls were made following the manufacturer protocol.

#### Sequencing

Sequencing was performed on the MGI DNBSEQ-G400, run Single Read 100 bp on a Large Flow Cell FCL SE100 (MGI). Datas were demultiplexed with BasecallLite v1.5.0.323.

#### Data analysis

Data analysis was performed by Guillaume Marcy and Maxime Lepetit (Labex Cortex Bioinformatics Platform).

Reads were processed to remove MGI adapter and low-quality sequences (Phred quality score below 20) using fastp version 0.24.0. Reads mapping and transcript expression quantification was performed onto the WBcel235 assembly (Ensemb release 113) using Salmon version 1.10.3 with Selective Alignment Full (SAF) approach, and converted to gene expression matrices with tximport R package version 1-26.1.

#### DEG analysis

DEG analysis was performed using the DESeq2 (v1.46) package in R (v4.4.0). Raw read counts were used as input, and genes with very low counts <10 across all samples were filtered out prior to analysis. DESeq2 uses the median of ratios method. Pairwise differential expression testing was carried out using the Wald test, and resulting p-values were adjusted for multiple testing using the Benjamini– Hochberg false discovery rate (FDR) procedure. Genes with an adjusted *p-value* < 0. 1 and an absolute log2 fold change ≥ 1 were considered significantly differentially expressed.

#### Sample age estimation

Analyses were performed in R (v4.5.0).

Sample age was inferred from TPM data with RAPToR (v1.2.0) (Bulteau & Francesconi, 2022).

L4 and YA samples were staged with the Cel_larv_YA reference (data from (Meeuse et al., 2020) from the associated data-package wormRef (v0.5.0). D7 samples were beyond the span of this reference, and staged on the Cel_aging_1 reference (data from GEO [byrneUnpublished]).

To confirm the D7 sample age estimates and get a common staging reference with earlier samples, we also staged YA and D7 samples with another aging reference (data from [hastings2020] and building method described in [bulteau2023thesis]), hereafter “hastings2020 reference”.

We subtracted the mutant and wild-type (WT) age estimates for each replicate and time point. Considering mutant and WT samples were collected at identical chronological times within each replicate, observed differences in biological (inferred) age should be caused by the genetic background.

Ages estimates of D7 samples (on the hastings2020 reference) were scaled to the time of the Cel_larv_YA reference using the YA samples (staged on both) to infer the linear relationship with a linear model (lm() R function).

## 10. Statistical analyses

Statistical analyses were performed using R (version 4.2.3) or Prism 10 (Graphpad). For all tests, compared samples were considered different when statistical test gave an adjusted *p*-value <0.05 (\**p* < 0.05; \*\**p* < 0.01; \*\*\**p* < 0.001), ns: non-significant.

## Supplementary figure legends

**Fig. S1.**
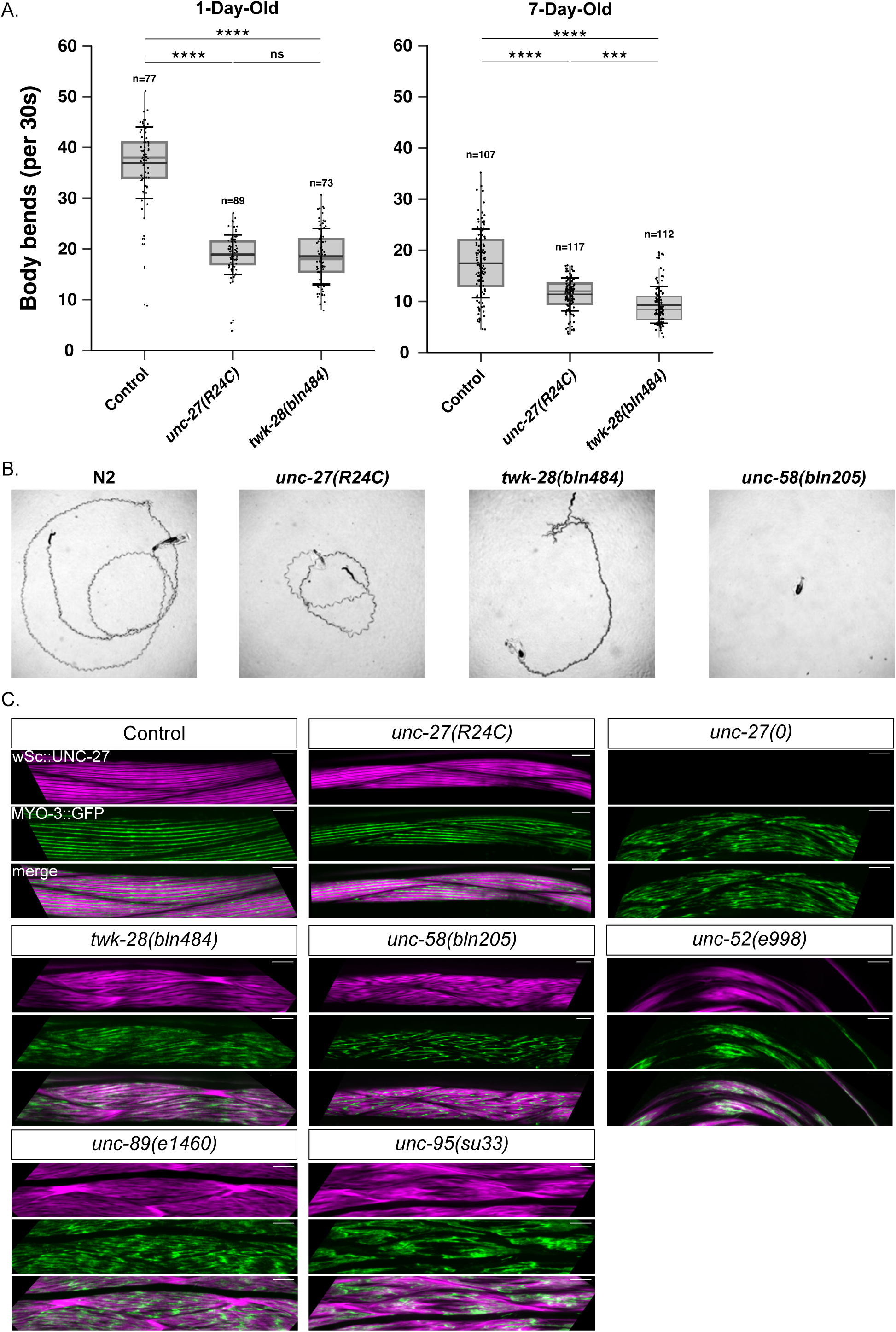
Comparison of the motility and sarcomeric organization of the body wall muscles in mutants used in. Figure 3. (A) Analysis of body bend frequency in 1 and 7-day-old control, *unc-27(R24C)*, and *twk-28(bln484)* mutants. The number of animals per group is indicated above the bars (pooled from three experiments). Boxes represent the interquartile range; gray lines indicate medians; black lines indicate mean; whiskers represent SD. Statistical significance was assessed by Kruskal-Wallis with Dunn’s post hoc test (FDR adjusted). (B) Representative images of movement traces of control, *unc-27(R24C)*, *twk-28(bln484)* and *unc-58(bln205)* on seeded NGM plates 5 minutes after worms were placed on the plates. (C) Representative images of myofilaments in young adult mutants of K2P channels and sarcomeric compartments. Thin and thick filaments are visualized thanks to wScarlet::UNC-27 and GFP::MYO-3, respectively. Scale bar = 10 µm.

**Fig. S2.**
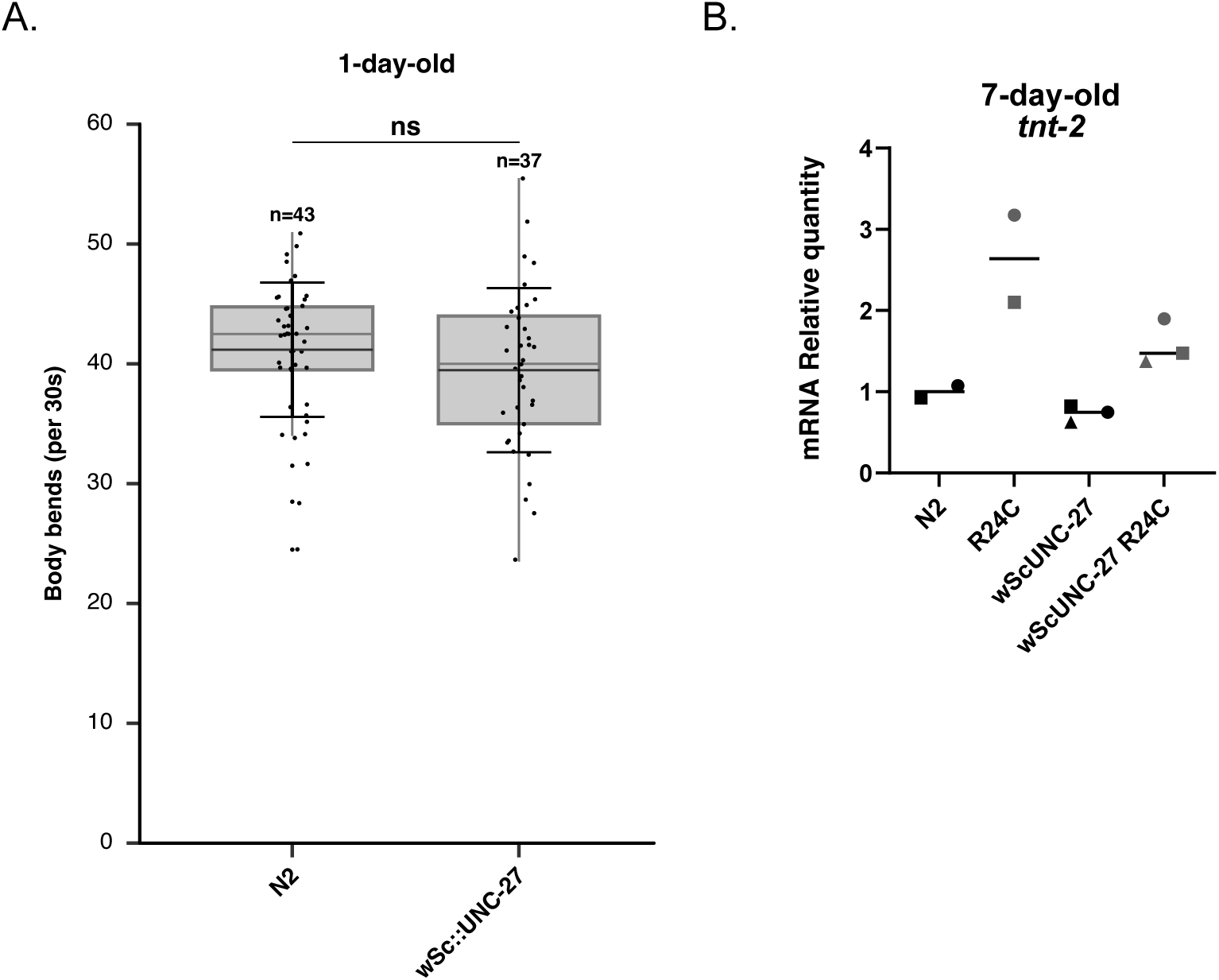
Validation of the functionality of the wScarlet-tagged version of *unc-27* in regulating worm motility and gene expression. (A) Analysis of body bend frequency in N2 and wScarlet::UNC-27 worms at the 1-day-old adult stage. Each dot represents an individual animal (n); boxes indicate the interquartile range; grey horizontal lines represent the median; and brackets denote the standard deviation. (B) Relative *tnt-2* mRNA levels in 7-day-old worms, comparing R24C and control worms in both untagged and wScarlet-tagged backgrounds. Each dot represents a biological replicate. Statistical significance for (A) was determined by a Wilcoxon Mann-Whitney test; ns, non-significant. No statistical analysis was performed for (B) due to the low number of biological samples.

